# Somatic DIS3 mutations in Multiple Myeloma arise early in clonal evolution, but are later counterselected due to toxicity

**DOI:** 10.1101/2023.07.27.550471

**Authors:** Tomasz M. Kuliński, Olga Gewartowska, Seweryn Mroczek, Marcin Szpila, Katarzyna Sałas, Vladyslava Liudkovska, Andrzej Dziembowski

## Abstract

DIS3 encodes an essential ribonucleic subunit of the nuclear exosome complex, responsible for degrading RNA in the nucleus. Somatic DIS3 mutations drive translocations in B cells, leading to multiple myeloma (MM). Clinical data analysis reveals that 42% of DIS3 mutations occur at three recurrent residues (D479, D488, and R780). These mutations, deactivating DIS3 exonucleolytic activity, are never homozygous, often appearing as minor subclones in advanced MM. Surprisingly, mutant DIS3 alleles undergo loss-of-heterozygosity, correlating with frequent del(13q) encompassing DIS3. Overexpression of wild-type DIS3 enhances growth and viability of DIS3-mutated MM cells, while CRISPR-mediated knock-out of the mutant variant, followed by longitudinal co-culture, replicates its elimination through counterselection, observed in the natural course of the disease. In mice, the heterozygous DIS3 *D479* mutation is embryolethal, confirming its dominant toxic effects. Transcriptome analysis of patients and cell lines reveals specific transcriptional signatures of DIS3 mutations with accumulation of non-coding unstable RNA species and including secondary indications of decreased proliferation. All these signatures are reversible upon mutant DIS3 loss-of-heterozygosity. DIS3 is an intriguing hit-and-run oncogene that drives MM, but is subsequently eliminated during clonal evolution.

## INTRODUCTION

Multiple myeloma (MM) is a neoplasm of plasma cells that predominantly affects elderly patients^1,2^. Clonal proliferation of MM cells in a bone marrow microenvironment leads to hematological and disseminated bone lesions. Moreover, high levels of monoclonal protein result in renal impairment or amyloidosis. MM accounts for 10% of all hematopoietic neoplasms, and its incidence has considerably increased over the past decades, making it a growing public health concern^3,4^.

The genetic disparateness of MM is a well-established phenomenon that contributes to its clinical complexity and heterogeneity^5,6^. Different patients exhibit a wide range of somatic genetic abnormalities that contribute to the development and progression of the disease. MM evolves from premalignant states such as monoclonal gammopathy of undetermined significance (MGUS) and smoldering multiple myeloma (SMM) through primary genetic events^6^. The presence of those genetic alterations classifies all MM cases into two groups: hyperdiploid MM, characterized by the presence of several chromosomal duplications and a more favorable clinical prognosis, and non-hyperdiploid (non-HRD), driven by chromosomal translocations affecting the immunoglobulin heavy chain locus^1^. Translocations involving this highly transcribed region, which contains strong enhancers, such as t(4;14), lead to the overexpression of oncogenes, including FGFR3, MMSET (NSD2), CCND3, CCND1, and MAF^1,6,7^. Deletions of chromosomal fragments, like del(13q), are also frequently observed, especially in non-HRD MM^8^. However, their role in disease progression is less well-defined^9,10^. Mutations or chromosomal abnormalities detected at the MGUS stage are considered primary events involved in tumor development. In contrast, events present in the later MM stages, but absent in MGUS, are considered secondary events leading to tumor progression. These secondary events may include acquired point mutations of importance, copy number abnormalities, and DNA hypomethylation. Interestingly, even though MM clonal evolution is intensely studied, the significance of loss-of-heterozygosity (LOH), one of the hallmarks of tumor evolution, is often overlooked.

Several somatic mutations in protein-coding genes have been described in MM, including the inactivation of tumor suppressor genes TP53 and RB1, as well as mutations in well-described oncogenes such as N-RAS and K-RAS^5,11,12^. Strikingly, mutations of two genes involved in RNA metabolism, TENT5C (FAM46C) and DIS3, are specific for MM5,^11,13,14^.

*DIS3,* located on chromosome 13, encodes a catalytic subunit of the primary eukaryotic nuclear ribonuclease exosome complex involved in the processing of stable RNA species, such as ribosomal RNA (rRNA), and clearing pervasive transcription products, such as promoter upstream transcripts (PROMPTs) and enhancer RNA (eRNA)^15–17^. DIS3 has two catalytic activities: the major exoribonucleolytic activity depending on the RNB domain and the endonucleolytic one arising from the PIN domain^18^. MM-specific DIS3 mutations are enriched in the RNB domain, inactivating or inhibiting the exoribonucleolytic activity of the enzyme. Therefore, they were assumed to alter posttranscriptional gene expression regulatory programs, leading to enhanced cell proliferation, specifically in the B-cell lineage^19–21^. This was even though DIS3 is an essential protein across species and cell types^16,22,23^. Moreover, even estimations of the stage of oncogenesis at which DIS3 plays a causal role are also conflicting^24–26^. Although mutations in protein-coding genes in MM are considered secondary events, DIS3 is mutated in MGUS, at the earliest stage of plasma cell dyscrasias^25^.

Our recent study has shown that mutation in DIS3 leads to an oncogenic transformation at the onset of myeloma progression^27^. DIS3 drives oncogenesis by promoting translocations involving the immunoglobulin heavy locus (IGH), which are well-known early drivers of myeloma. This occurs by targeting activation-induced cytidine deaminase (AID) to chromatin due to the stalling of DIS3 on native transcripts. AID plays a crucial role in the immune response by diversifying immunoglobulin repertoire through somatic hypermutation (SHM) and class-switch recombination (CSR). However, it must be tightly controlled to minimize off-target activity^28,29^. DIS3 and the nuclear exosome have been shown to play a role in AID regulation^30–33^. Thus, the MM DIS3 alleles promote AID promiscuous activity directly through its attraction to chromatin.

This study aims to elucidate the role of DIS3 mutated alleles in clinically manifesting MM and concentrates on its effects on cell physiology. Through comprehensive analysis of clinical genomic data, we identified recurrent DIS3 mutant alleles at residues: D479, D488, and R780, which, during MM progression, behave in a very unusual way. Recurrent DIS3 alleles are consistently heterozygous or represent minor subclones. Notably, the mutated variant, not the wild-type (WT) one, undergoes LOH during MM development. Generally, all MM-associated DIS3 variants perturb RNA metabolism, leading to the accumulation of natural exosome substrates in MM cells. Counterintuitively, these mutations adversely affect cell survival, resulting in reduced proliferation of MM cell lines. Recurrent MM DIS3 alleles are embryonically lethal, even in the heterozygous state. By longitudinally co-culturing patient-derived cells with CRISPR-induced KO of the mutant variant, we replicate *in vitro* the counterselection of DIS3 MM alleles during disease’s clonal evolution. In sum, we describe DIS3 as a new model oncogene that arises early to drive oncogenesis but is then counterselected.

## RESULTS

### Identification of recurrent DIS3 mutations D479, D488, and R780

To assess the spectrum of DIS3 mutations in MM, we took advantage of the ongoing Multiple Myeloma Research Foundation CoMMpass study. Whole-exome sequencing data analyses revealed DIS3 missense mutations in 14% of cases (132 of 993; Figure 1A, B, Figure EV1, Table EV1 and EV2). Consistent with previous findings, observed amino acid substitutions within DIS3 were strongly enriched within its exoribonucleolytic RNB domain (Figure 1B)^11,16^. Notably, 33% of somatic variants were detected at three recurrent locations: D479, D488, and R780. These mutations completely inactivate RNB domain activity^16^. Although distant in the protein sequence, the residues are brought in close proximity in the active site of DIS3^34,35^, which structure was previously resolved by electron microscopy^34^. D479 and D488 are catalytic residues, whereas R780 positions the substrate for catalysis (Figure 1C)^16^. Other identified mutations were also enriched in highly conserved residues of the RNB domain and sites that interact with RNA substrates (Figure 1D). At the same time, no mutations predicted to induce dissociation of the enzyme from the exosome complex have been found (Figure 1D), showing that gain rather than loss of function mutations are associated with MM. These mutations are predicted to have a high probability of pathogenicity based on the PON-P2 algorithm^36^, altogether suggesting a similar mode of action as recurrent mutations but with varying potency (Figure 1E). In addition to somatic mutations, we identified germline DIS3 variants in the CoMMpass study, although they are predicted to be less pathogenic than somatic ones (Figure 1F, G). Some variants were found in a homozygous state, suggesting their weaker impact on cell physiology but still showing enrichment compared with the general population (Table EV1, EV2). Germline variants were previously implicated as the basis of familial MM cases^37^.

**Figure 1.**
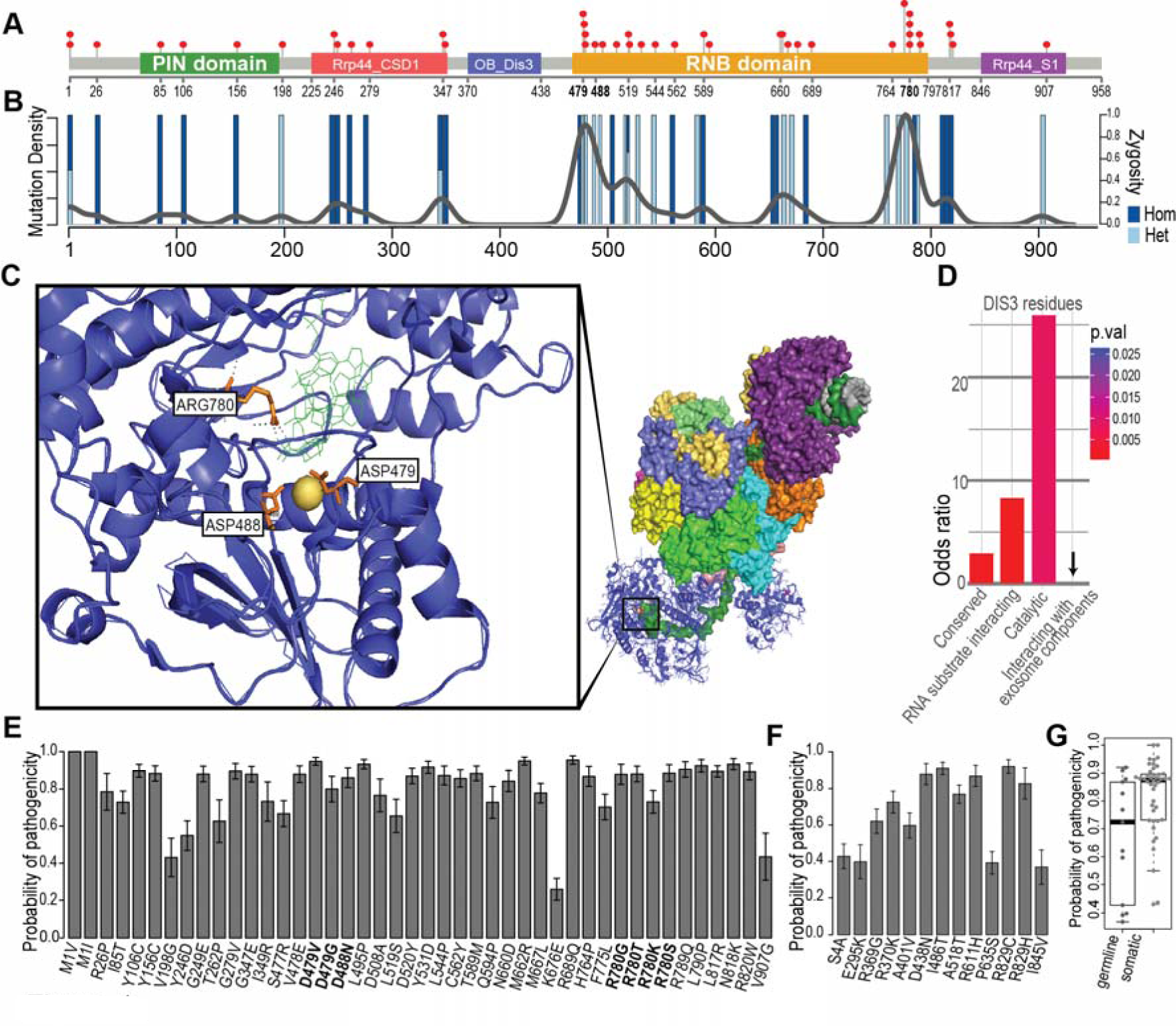
Recurrent MM DIS3 mutations (D479, D488, and R780) with possible dominant-negative effects. (A) Distribution of somatic DIS3 mutations in 901 MM cases in the CoMMpass study. Lollipops indicate the mutation location relative to the DIS3 domain structure. (B) Mutation density and zygosity frequency are plotted below. (C) The recurrently mutated residues D479, D488, and R780 are located in the DIS3 active site, affecting the coordination of the catalytic Mg^2+^ ion or positioning of the RNA substrate backbone for catalysis. (D) Somatic mutations are significantly enriched in catalytic residues, highly conserved residues, and residues that interact with the RNA substrate (Fisher’s Exact Test). None of the sequenced mutations affected residues responsible for the interaction between DIS3 and other components of the exosome complex. (E) Prediction of pathogenicity of somatic DIS3 variants by PON-P2. Somatic mutations are generally highly likely to be pathogenic. Error bars represent the standard deviation. (F) PON-P2 prediction of the pathogenicity of germline DIS3 variants. (G) Comparison of PON-P2 prediction of the pathogenicity of germline and somatic DIS3 variants.

Notably, no nonsense DIS3 mutations were found in MM patients (Table EV1). At the same time, recurrent mutations are exclusively heterozygous, as by quantification of the variant alleles frequency (VAF) (Figure 1B, Table EV1). All this suggests that DIS3 mutations result in a gain of function and/or have dominant-negative effects, indicating that depletion-based experiments are not proper models of DIS3-dependent cancerogenesis.

### Recurrent DIS3 alleles are counterselected at late stages of disease progression

Next, we focused on the recurrent alleles we identified (D479, D488, and R780), which are expected to have the strongest cancerogenic effects. Notably, three subjects of the CoMMpass study bearing DIS3 mutations had their MM samples sequenced at two stages of disease progression. In all cases, these were recurrent alleles (two D488N and one R780K). VAF analysis revealed that, surprisingly, mutant alleles in all three of those cases were lost or became minor subclones during MM evolution (Figure 2A). This loss-of-heterozygosity contradicts the previously proposed tumor suppressor mode of action of DIS3^16,19,23^.

**Figure 2.**
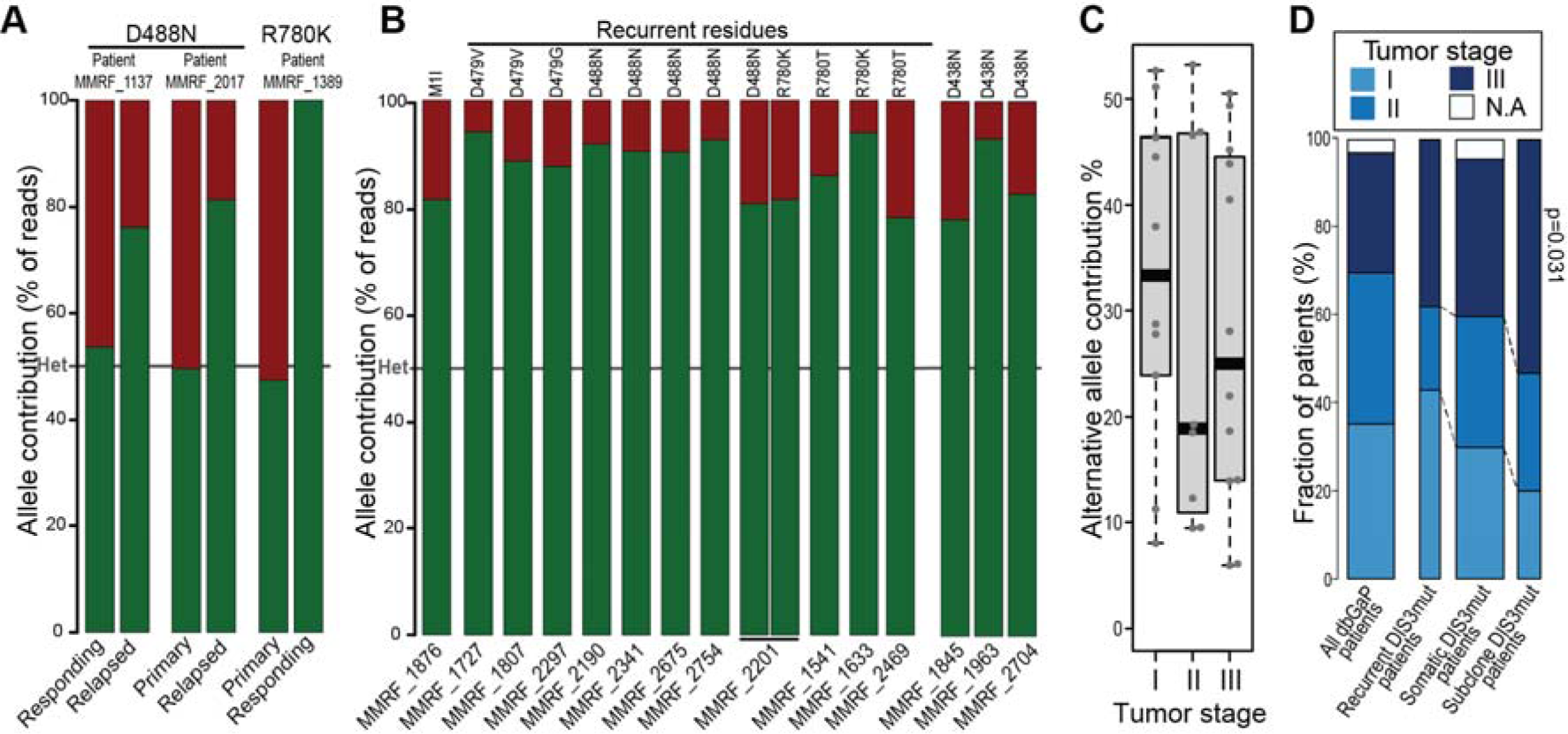
DIS3 loss-of-heterozygosity with the retention of the WT allele is common in MM patients. (A) Contribution of recurrent DIS3 mutations in biopsies of patients in the CoMMpass study sequenced at two stages of MM progression based on whole exome sequencing (WES). (B) The contribution of WT and mutant DIS3 alleles in MM patients with recurrent DIS3 mutations present as a minor subclone, based on WES. (C) The contribution of recurrent mutant DIS3 alleles in MM patients sequenced at different MM stages. (D) Staging of MM for cases showing at their first sample sequenced DIS3 mutant allele as a minor subclone compared with the distribution in all patients. The group of patients with a MM DIS3 variant present as a minor subclone is significantly enriched in the later stages of the disease (Fisher’s Exact Test, *p* = 0.031).

The presence of DIS3 alleles as minor subclones prompted us to reanalyze the CoMMpass dataset. This revealed additional 12 MM subjects with DIS3 mutations present as a minor subclone at the time of their enrollment in the study (Figure 2B). When considered, recurrent variants accounted for 42% of all somatic DIS3 mutations. Patients with minor DIS3 subclones were found to be in the later stages of MM compared with the rest carrying DIS3 mutations. In contrast, subjects with heterozygous states of recurrent DIS3 alleles were prevalent in the first stage of MM (Figure 2C, D), strongly indicating that the observed subclonality is a consequence of counterselection during disease progression. At least for a fraction of cases, the LOH of the recurrent DIS3 allele is caused by del(13q) encompassing the DIS3 loci^21^.

### DIS3 mutations are toxic to cell physiology affecting proliferation

Knowing that the depletion of DIS3 does not mimic the pathophysiology of MM, we first searched for homozygous mutations in established MM cell lines^38^ to develop an appropriate cellular model. No single case of homozygous recurrent mutation was found at the DIS3 locus among the genomes of 70 MM lines sequenced. However, we identified a homozygous A507P mutation in the DIS3 RNB domain in the PE2 cell line, which inhibits but does not fully inactivate the enzyme^39^ (Figure EV2A).

To examine the potential oncogenic contribution of the DIS3 mutation in PE2 cells, we introduced WT DIS3 using a lentiviral vector. The reintroduction of WT DIS3 increased the growth rate of PE2 cells compared to control green fluorescent protein (GFP)-expressing cells, reducing the doubling time by 21% (Figure 3A). The increase in the proliferation rate was also clearly visible in CellTrace assay operating on the principle of fluorescence signal dilution during cell division (Figure 3B). At the same time, overexpression of WT DIS3 significantly improved PE2 cell viability (Figure 3C, Figure EV2B).

**Figure 3.**
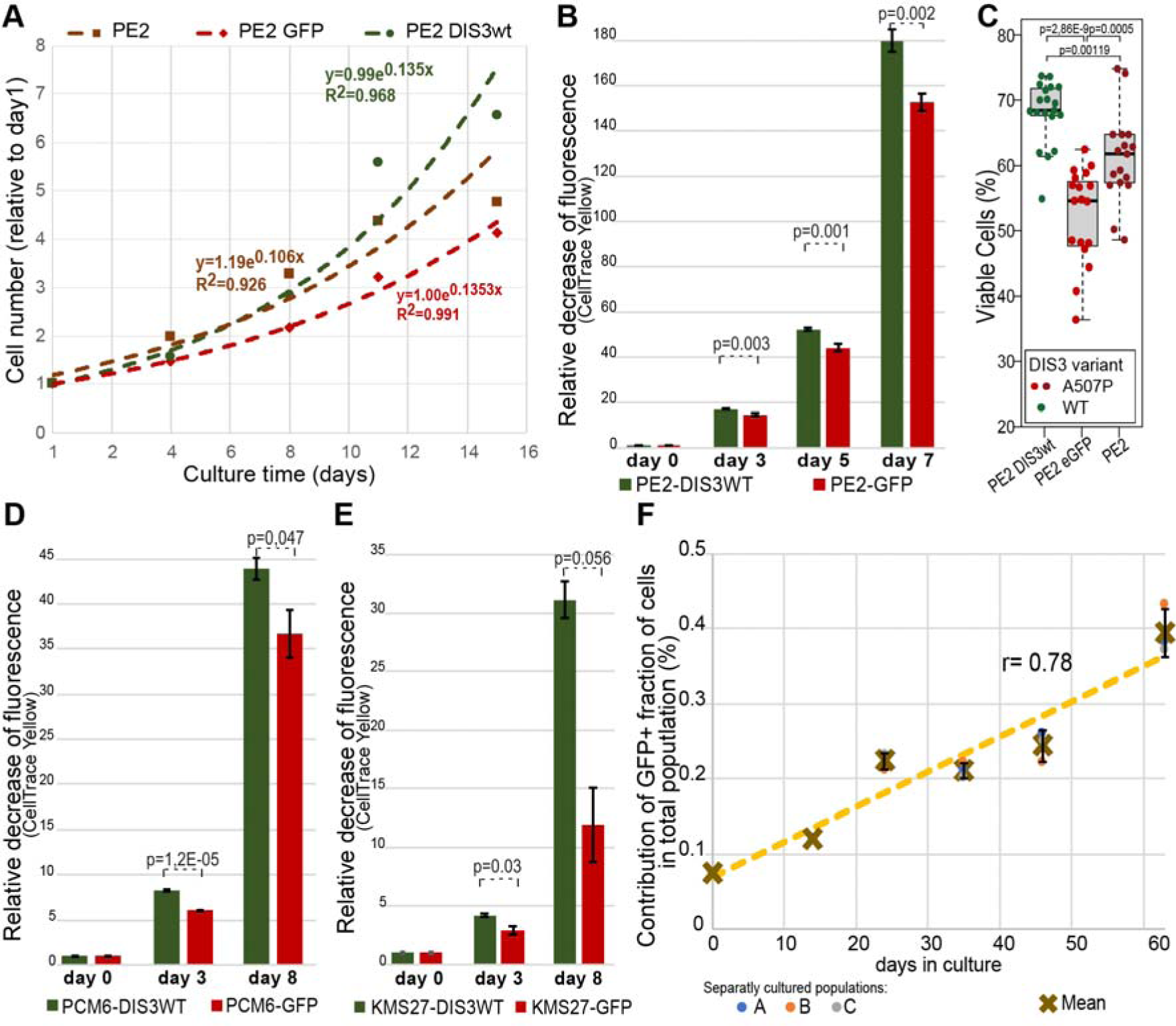
DIS3 RNB mutant protein variants decrease proliferation rate and viability of MM cell lines. (A-B) the proliferation rate of PE2 cell line with homozygous DIS3^A507P^ after overexpression of WT DIS3 measured by cell counting (A) and CellTrace assay based on fluorescence signal dilution during cell division (n=3) (B). (C) Cell viability of PE2 cells with DIS3 WT overexpression and GFP as a control, measured by fluorescent dead cell discriminator and flow cytometry (n=3). (D-E) The proliferation rate of cell lines measured with CellTrace assay in heterozygous recurrent DIS3 mutations: KMS-27 (D488N) (D) and PCM6 (R780K) (E) and after overexpression of WT DIS3. In 3A-E, error bars represent standard error (performed in triplicates co-stained and cultured in parallel for each cell line). (F) Long-term co-culture of KMS-27 MM cells with their DIS3 D488N KO shows a gradual increase of hemizygous WT cells contribution in the population as measured by GFP fluorescence.

Next, we examined MM cell lines with heterozygous recurrent DIS3 mutations, namely KMS-27 (harboring D488N) and PCM6 (harboring R780K). Similar to the PE2 cell line, the overexpression of WT DIS3 enhanced the proliferation rate of these cell lines (Figure 3D-E). To mimic the counterselection of the mutant allele observed in patients, using cell culture, we transfected the KMS-27 cell line, which is heterozygous for the D488N variant, with CRISPR/Cas9 constructs, knocking out the mutated DIS3 variant by truncating it in the second exon and incorporating a fluorescent GFP marker. Longitudinal co-culturing of KMS-27 cells with their DIS D488N knockout revealed an increase in the proportion of cells expressing GFP in the population, indicating a selective advantage of the hemizygous DIS3 WT genotype (Figure 3F and Supplementary Figure 2C).

In sum, MM-related DIS3 variants impede cell survival and proliferation, even in MM cells, generating a selective pressure for the LOH of the mutated allele.

### DIS3 mutations lead to specific transcriptomic signatures which are reversible upon LOH

To find dysregulated cancer-related transcripts in DIS3 mutated cells, we analyzed RNA sequencing (RNA-seq) data from the CoMMpass study and compared transcriptomes of patients with and without DIS3 mutations. DIS3 mutations led to the accumulation of expected DIS3 substrates PROMPTs (Figure 4A, B) and rRNA processing byproducts (Figure EV3A). Generally, long non-coding RNA (lncRNA) represented the majority of transcripts accumulating in patients with defective DIS3 (Figure 4A-C, Figure EV3C and Data S1, S2). These transcripts are predominately nuclear-localized and are degraded by the nuclear exosome^17^. There is a substantial overlap in accumulating RNA species between recurrent and non-recurrent MM DIS3 alleles (Figure EV3B). In contrast to ncRNAs, dysregulated protein-coding transcripts constitute only 9% and 18% of the most significantly accumulating transcripts in recurrent and non-recurrent MM DIS3 mutant patients, respectively (Figure 4C). In both patients without mutated DIS3 and PE2 cell line overexpressing WT DIS3, the accumulation of mRNAs immunoglobin mRNAs is observed, which may be a consequence of their high transcriptional and nuclear transport rate, making them least affected by dysfunction of the nuclear exosome. Immunoglobulin mRNAs are not expected to impact oncogenesis (Dataset EV1, EV2).

**Figure 4.**
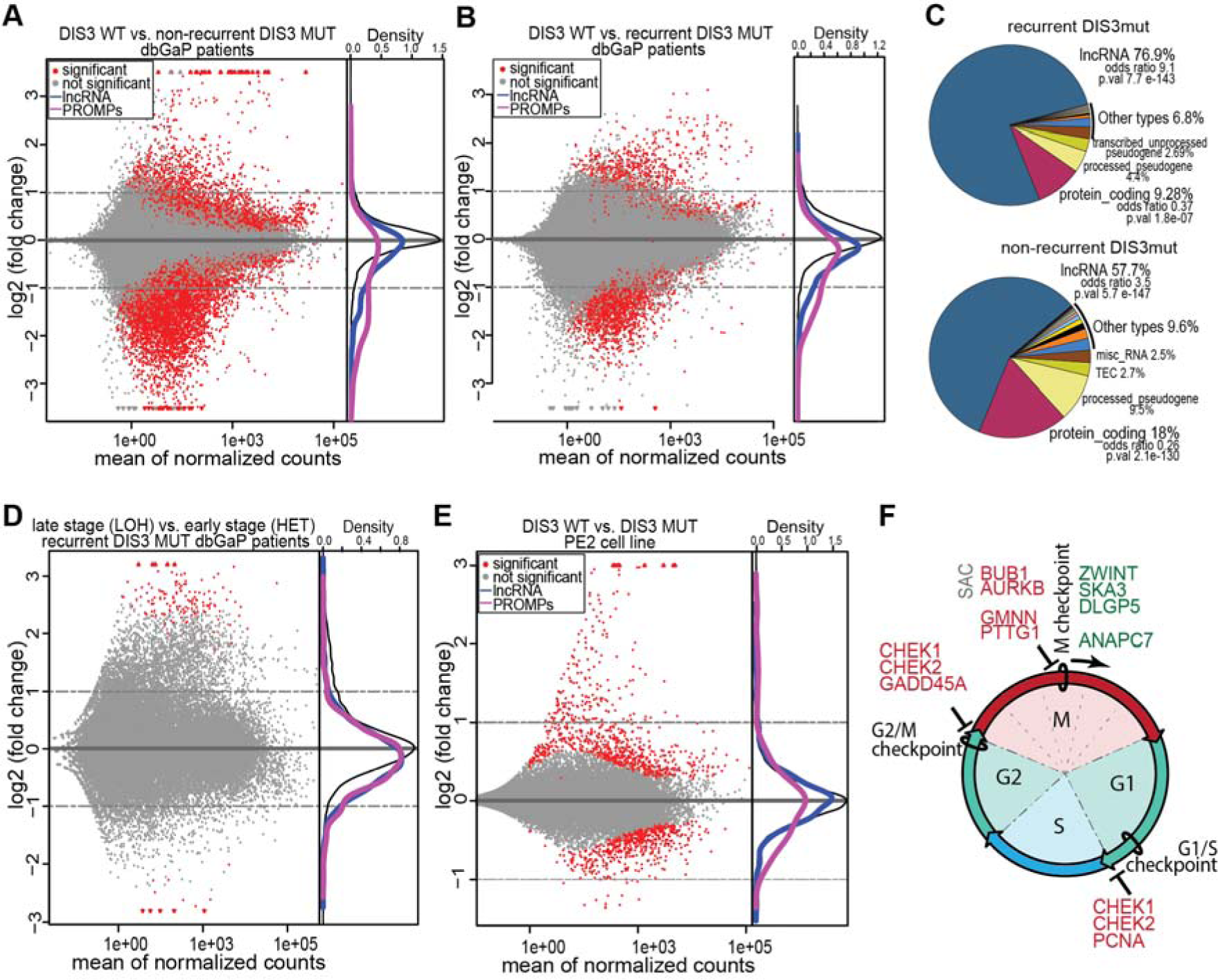
Transcriptional signatures of MM DIS3 alleles. (A, B) MA plots illustrating the differences between transcriptomes of patients with non-recurrent and recurrent DIS3 mutations compared to the control group with WT DIS3. Density plots in the panels on the right show the global accumulation of PROMPTs and lncRNAs in patients with an MM-specific DIS3 mutation. (C) Overlap of most significantly dysregulated genes in DIS3 recurrent mutations and non-recurrent DIS3 somatic mutations. (D) MA plots displaying changes in the transcriptome of LOH patients upon the loss of the recurrent DIS3 mutant allele. Density plots in the panels on the right show the global accumulation of PROMPTs and lncRNAs in patients with an MM-specific DIS3 mutation. The comparison is between 2 early samples and 3 LOH samples. (E) MA plot showing changes in the PE2 transcriptome upon the reintroduction of WT DIS3. Density plots in the panels on the right show the global downregulation of PROMPT and lncRNA in cells with reintroduced DIS3 WT. (F) Schematic diagram depicting the dysregulation of transcripts involved in progression through mitosis during checkpoints, accumulated in DIS3 mutant (red) and WT DIS3 (green) cells.

Comparing the transcriptomes of patients with MM-specific recurrent DIS3 mutant alleles before and after the LOH revealed a reversal of the nuclear non-protein-coding transcript accumulation phenotype (Figure 4D, Figure EV3D). Notably, specific changes in the expression patterns of protein-coding genes become evident for LOH patients. The depletion of MM-specific DIS3 variants downregulates genes involved in cell transition through mitosis (Figure EV3E, Dataset EV3), consistent with the potential role of DIS3 in kinetochore formation described for its yeast ortholog^22,23^ (Figure EV3E). Still, it contrasts with the previously postulated positive effect of DIS3 mutations on proliferation in MM^19,20^. In a cellular model, the overexpression of WT DIS3 in the PE2 MM cell line reduces the accumulation of DIS3 substrates (Figure 4E, Figure EV3F). At the same time, it leads to the downregulation of DNA stress response genes involved in the base excision repair (BER) and DNA mismatch repair (MMR) pathways, which participate in class switch recombination during B cell activation^40–42^ (Figure EV3H, Dataset EV4). Similarly, in the LOH patient sample analysis, genes involved in the cell transition through mitosis are dysregulated (Figure 4F, Figure EV3E, Dataset EV4). Cells with the overrepresentation of the mutant DIS3 allele upregulate genes that inhibit the transition through checkpoints (CHEK1, CHEK2, PCNA, GADD45A, GMNN, and PTTG1) and the spindle assembly complex (SAC; Figure 4F). Conversely, cells with the overrepresentation of WT DIS3 exhibit the upregulation of genes responsible for the transition through the spindle checkpoint, such as components of the anaphase-promoting complex (APC/C; Figure 4F).

Altogether, our transcriptomic analysis of MM patients and MM cell lines revealed detectable transcriptional signatures associated with DIS3 mutations, which are unlikely to drive oncogenesis. Notably, transcriptomic effects are reversible since they disappear as a consequence of LOH of the DIS3 mutant allele.

### Recurrent DIS3 mutations induce mitotic arrest at G1 and G2 checkpoints and are embryolethal in mice at heterozygous state

As described above, in agreement with the negative effect of DIS3 mutations on proliferation, transcriptomic analysis of patient samples and MM cell lines revealed an association of DIS3 mutation with the deregulation of several genes involved in mitosis and DNA replication (Figure 4F, Figure EV3G-H), which is also evident in pathway analysis (Figure EV3I). We used an established cellular model based on HEK293 cells to dissect the nature of these effects further^16^. This model allows for the inducible replacement of an endogenous allele with a mutant variant harboring DIS3 catalytic mutation, equivalent to mutations observed in patients. Indeed, DIS3 mutations resulted in a strong inhibition of proliferation (Figure EV4A and Figure EV4B), with mitosis being blocked at both G1 and G2 checkpoints (Figure 5A). The character of this mitotic dysfunction could correspond to the changes observed in the transcriptome analysis (Figure 4F, Figure EV3I), particularly the activation of the p53 pathway (Figure EV3I). Simple DIS3 depletion was also recently shown to affect the proliferation of MM cell lines^43^.

**Figure 5.**
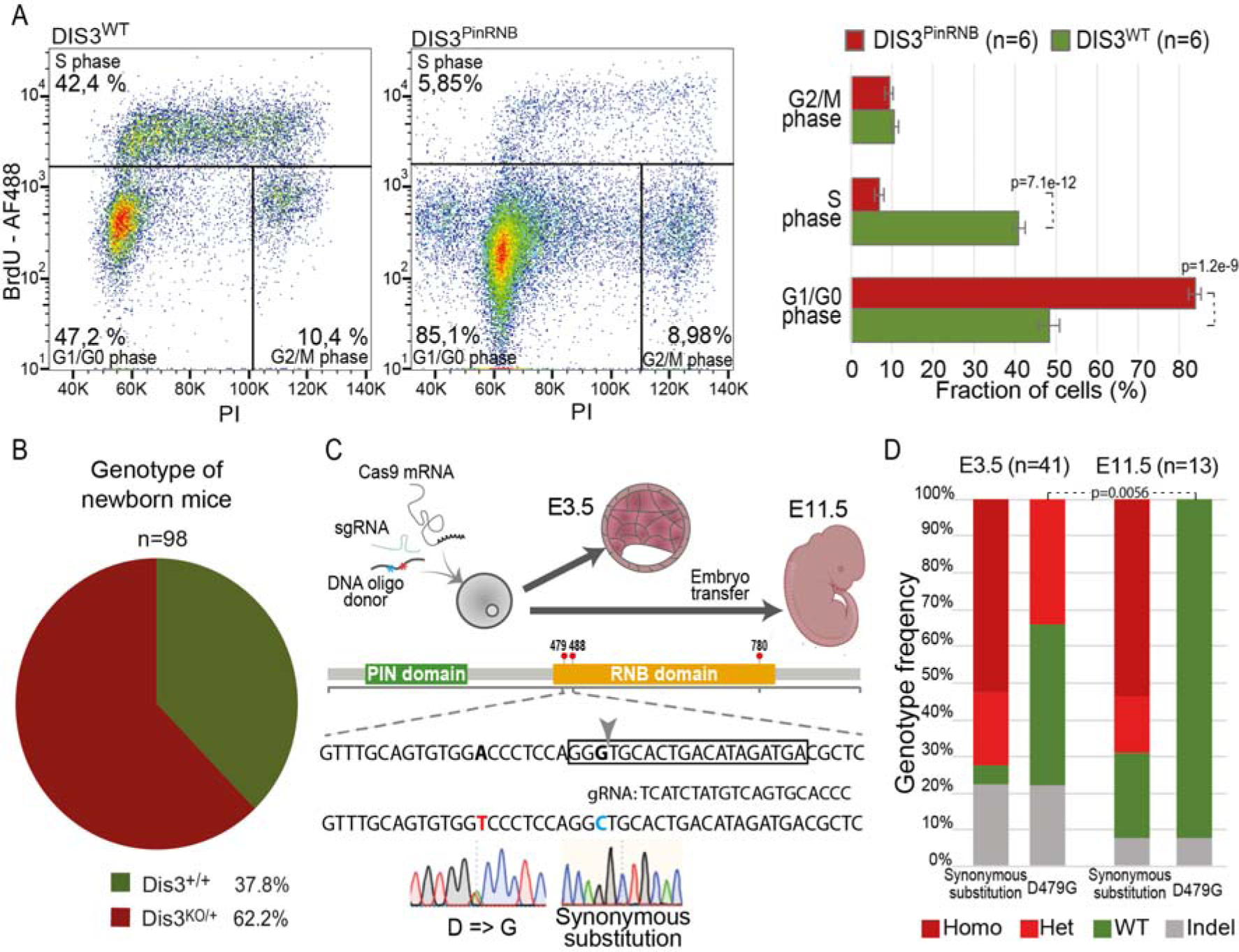
Strong DIS3 mutations lead to mitosis arrest and are embryonic lethal at early stages in mice. (A) BrdU/PI staining visualizes mitotic arrest at G1 and G2 checkpoints after induced expression of a D487R DIS3 catalytic mutant. (B) Frequencies of indicated genotypes in offspring of Dis3^KO/+^ x Dis3^KO/+^ matings show reviling viability of heterozygous KO. (C) Schematic illustration of CRISPR/Cas9 strategy of introducing recurrent mutations in mice. Position of the single-guide RNA (sgRNA) target is indicated with a box and the introduced substitutions and the “cut” position are marked. (D) The frequency of genotype occurrence for co-introduced D479G and a synonymous substitution at E3.5 and E11.5 embryonic development stage. Heterozygous recurrent D479G mutants show embryonic lethality at an early stage. (Homo - homozygous, Het - heterozygous, Indel – an insertion or deletion on both alleles).

**Figure 6.**
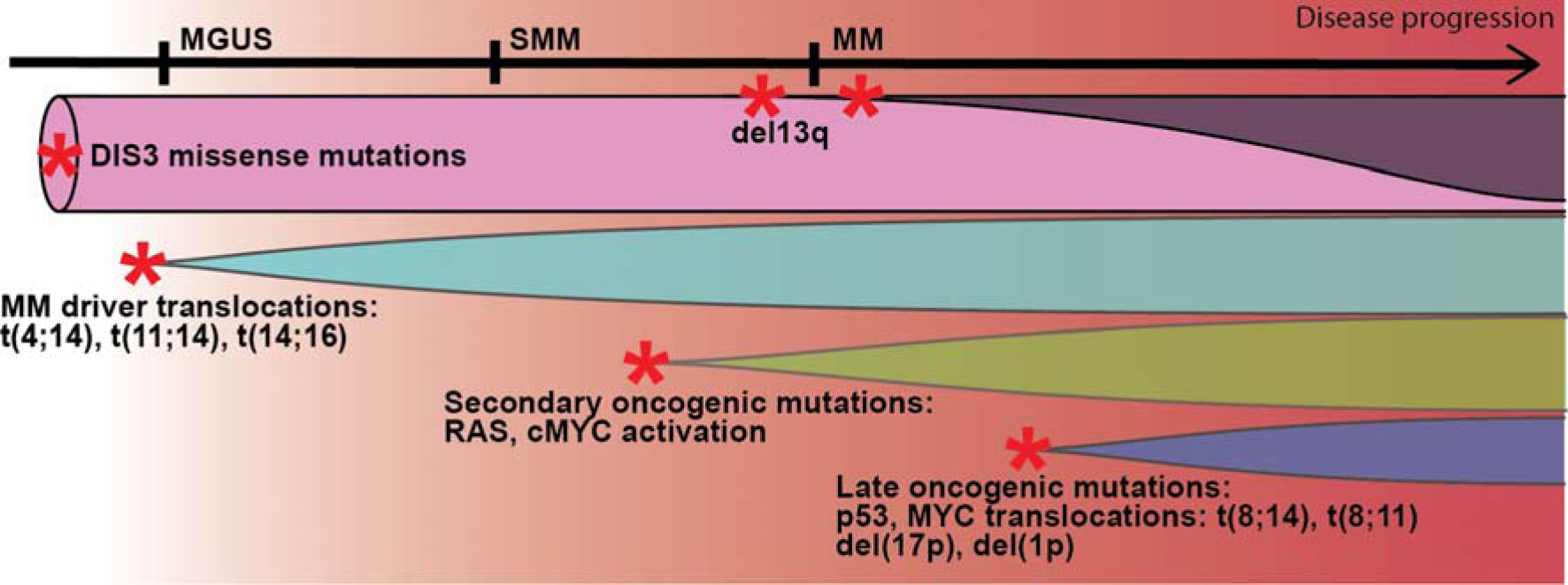
Proposed involvement of DIS3 in clonal evolution during disease progression of plasma cell dyscrasias. DIS3 MM-specific missense mutations arise in the early disease stage of MM, likely in pre-germinal center cells. MM-specific DIS3 variants contribute to genome destabilization, thereby increasing the incidence of oncogenic translocations within the early window of opportunity during naive or memory B cell activation. As the disease progresses to later stages, the mutation becomes dispensable for the oncogenic state and instead becomes a burden for the cell, leading to its loss, such as through the deletion of the arm of chromosome 13q containing it. The 13q deletion can occur at any stage of the progression of plasma cell dyscrasias. However, given/considering the proliferative phenotype, the strongest selective pressure would be present at the MM stage when cell expansion is the highest.

Considering the predominantly negative impact of MM DIS3 alleles on cell proliferation and survival, we aimed to dissect their potential oncogenic effects using an animal model. First, we introduced a knockout of the DIS3 gene by introducing a frameshift insertion in exon 3 using CRISPR/Cas9 methodology. DIS3 KO leads to embryolethality when homozygous (Figure 5B). In a heterozygous state, the knockout allele does not produce any visible phenotype in mice bred under standard conditions (Figure 5B and Figure EV4C). Therefore, we attempted to generate mouse lines harboring D479V, D488N, or R780T mutation using CRISPR/Cas9 methodology. We have failed in all cases, as no mice with the desired mutation were born. Instead, we obtained several mice with heterozygous indels causing frameshifts (equivalent to constitutive KO), suggesting that the dominant negative effect of recurrent DIS3 MM alleles leads to embryolethality. To examine the quantitative effect of these mutations, we focused on D479V and analyzed the embryos after microinjections (Figure 5C). Out of the 50 that were cultured, 41 progressed to the blastocyst stage and were genotyped. All blastocysts were successfully modified (Figure 5D), with the majority displaying indel mutations resulting in frameshifts. Among the blastocysts, 19,5% carried the heterozygotic catalytic mutation (D479V/WT), while none had the homozygotic catalytic mutation, even though homozygotic silent mutation was observed in almost all non-indel cases (Figure 5D). In parallel with the analysis of blastocysts, 50 two-cell embryos were transferred to four surrogate mothers. Three of the transfers resulted in pregnancies, and at E11.5 stage, we found 13 embryos and multiple implantation sites with resorptions. However, upon genotyping of these 13 embryos, none were found to harbor the D479V mutation in either a homozygous or heterozygous state, although most displayed accompanying silent mutations. Because a single allele knockout by frameshift indels does not affect the viability of E11.5 embryos or mice, D479V mutation has a clear dominant negative effect leading to early embryonic lethality.

## DISCUSSION

The role of DIS3 mutations identified in MM has remained enigmatic for years, as neither the mechanism by which they drive oncogenesis nor their evolution and direct effect on cell physiology have been well described. We recently described the mechanism of oncogenic transformation resulting from DIS3 mutations, which is based on the induction of activation-induced cytidine deaminase (AID)-dependent chromosomal driver translocations at the early stage of MM^27^. This study describes an unusual clonal evolution of MM driven by recurrent DIS3 alleles, eventually resulting in their counterselection. It focuses on recurrent mutations, which, as we show, represent at least 42% of all genetic alterations in DIS. Due to the LOH of mutated alleles, the exact frequency of DIS3 somatic mutations in MM is difficult to estimate. However, the number is expected to be even higher.

The majority of previous experimental work on DIS3 in the context of MM employed KD or KO models^43–45^, even though DIS3 is an essential gene and only point missense mutations are found in MM. Analysis of DIS3 point substitutions, the absence of mutations that would result in the loss of the protein product or the dissociation of the exosome complex, together with their heterozygous state, suggest that they are dominant negative. Biochemical analysis of some of those mutations or similar ones has shown hampered enzymatic processivity of DIS3^16^. Such mutations stall the nuclear exosome on its natural RNA substrates^17^. MM DIS3 mutations, especially the recurrent ones (D479, D488, and R780), are expected to change the nuclear exosome complex into an RNA-binding protein rather than the active exoribonucleolytic enzyme. Moreover, recurrent alleles cause early embryonic lethality in heterozygous mice. In contrast, knockout alleles do not produce any visible phenotype when heterozygous. This is in concordance with the lack of phenotype of the conditional presumably hypomorphic Dis3-COIN allele in cultured B cells^32,44^. The nature of the described clinical DIS3 mutations supports the model explaining the exosome complex stalling on chromatin^27^, and underlining the need for using models reproducing properties of MM alleles when studying MM pathogenesis.

DIS3 mutations in MM patients are associated with pleiotropic changes at the transcriptome level, which previously were proposed to drive oncogenesis^46^. Differential RNA steady state analysis revealed predominantly an accumulation of non-protein coding transcripts, including byproducts of rRNA processing and nuclear lncRNAs, and a minor effect on the protein coding transcriptome. GO analysis of perturbations in signaling pathways demonstrates decreased mitotic potential of DIS3 mutated cells. Our data strongly support the interpretation that the oncogenic effect of DIS3 mutation cannot be attributed to transcriptome deregulation. Notably, transcriptomic effects are reversible as a consequence of LOH of the DIS3 mutant allele. This is accompanied by the progression of MM patients to later disease stages and an increase of the proliferative potential of MM cell lines cultured *in vitro*. Accumulation of nuclear unstable ncRNAs was also noticed by others, who, however, suggest that these play a positive role in oncogenesis^21^. The toxicity of MM DIS3 alleles in MM could be explained by the accumulation of RNA species that are typically swiftly cleared by the nuclear exosome, but the exact mechanism is yet to be revealed. The impaired transition through mitosis is expected to become a greater burden in the later malignant stages of plasma cell dyscrasias, during which cell proliferation significantly increases.

High-throughput DNA sequencing from MM patients reveals the presence of at least two compensatory mechanisms that enable the overcoming of the toxic effects caused by DIS3 alleles during MM evolution: frequent LOH of DIS3 alleles and acquisition of mutations that inactivate tumor-suppressive pathways and are pro-proliferative. The del(13q) chromosomal region includes the RB1 gene and Mir15a/Mir16-1 miRNA cluster^47,48^. The RB1 encodes a crucial tumor suppressor that safeguards the G1/S checkpoint, the haploinsufficiency of which is known in many cancers, including MM^49^. The Mir15a/Mir16-1 miRNA cluster also plays a tumor-suppressive role by negatively regulating mitosis^47^. Similarly, DIS3 mutations are known to significantly co-occur with 1q21.3 gain (including CKS1B), which positively regulates mitosis entry and the deactivation of TRAF2^5^. The significant co-occurrence of DIS3 mutations with RB1 or TRAF2 deactivation^5,47^ suggests that DIS3 mutations can additionally contribute to greater genomic drift in MM, affecting proliferation and, consequently, the selection of deactivating mutations in tumor suppressors. As we showed in a recent study, the MM-specific DIS3 variant, especially the more devastating recurrent mutations, are mechanistically linked with the co-occurrence of a t(4;14) translocation, which activates FGFR3, providing an oncogenic signal to enhance cell proliferation^27^. The pro-proliferative signal coupled with faulty cell divisions leads to mitotic stress and eventually causes the counterselection of the defective DIS3 allele. This likely explains the high level of co-existence of t(4;14) and del(13q) observed in MM patients^50,51^.

According to our data, DIS3 is a unique gene associated with cancer. DIS3 mutations drive carcinogenesis through mutagenic effects at an early window of opportunity during MM oncogenesis^27^. Genomic instability, known as one of the enabling characteristics of cancer^52^ in the case of MM happens in a short window of opportunity during B cell activation^27^. DIS3 recurrent alleles essential for initiation of the MM oncogenic transformation at pre-clinical stages become a burden for the ever more rapidly dividing cancer cells and are eliminated during the clonal evolution at later stages of diseases progression. There are examples of genes with contradicting effects on cancerogenesis, such as NOTCH, whose activity is known to have opposing properties from oncogenic to tumor suppressive, depending on the cancer type and the genetic background^53–55^. However, the MM DIS3 variant serves as a unique “hit-and-run” oncogene displaying both canacer-promoting, as well as, antiproliferative properties in a cancerous cell during the progression of the disease in a single patient. The example of MM recurrent DIS3 alleles clonal evolution implies the necessity to reconsider the common assumption that subclonal mutations must be late acquisitions by the tumor cells. With massive sequencing efforts for most cancer types and analyses of samples at several time points of disease progression, future studies could identify other oncogenes that behave similarly. Understanding this peculiar class of oncogenes, introducing vulnerabilities to cancer cells just as DIS3 mutations is essential for the development of individualized therapeutic strategies for the targeted treatment of cancer patients.

## Methods

### PE2 cell line

The Human Multiple Myeloma PE2 cell line was kindly provided by Prof. Michael Kuehl from the Center for Cancer Research, National Cancer Institute. Cells were cultured in RPMI 1640 ATCC’s modified medium (Invitrogen; A1049101) supplemented with 4% FBS (Invitrogen; 10270-106), penicillin/streptomycin (Sigma; P4083) and 2 ng/ml of recombinant human IL6 (PeproTech; 200-06) at density between 0.5-1 x 10^6^ cells/ml. Media were changed every 3 days. The *DIS3* gene was cloned from human cDNA into pcDNA^TM^5/FRT/TO vector (Invitrogen) using SLIC protocol as described^56^. To obtain SFFV (pLVX) lentiviral constructs, the *DIS3* inserts were generated by PCR using pcDNA-DIS3-TEV-eGFP as previously described^14^. Lentiviruses expressing GFP only were described previously^17^.

For proliferation rate assessment of PE2 cell lines, cells were seeded at the density of 0.5×10^6^ cells/ml and counted using a hemocytometer. Viability assessment was performed by staining cells with LIVE/DEAD™; Fixable Green Dead Cell Stain Kit (Invitrogen; L2310) according to the manufacturer’s instructions. Cells were analyzed on Attune NxT (Thermo Fisher Scientific) or CytoFLEX LX (Beckman Coulter) flow cytometers.

### KMS-27 and PCM6 MM cell lines

The KMS-27 Human Multiple Myeloma cell line baring a D488N DIS3 allele was acquired from the JCRB Cell Bank (Japanese Collection of Research Bioresources Cell Bank), whereas the PCM6 RIKEN BioResource Research Center, Koyadai Tsukuba-shi Ibaraki, JAPAN. KMS-27 cells were cultured in RPMI 1640 ATCC’s modified medium (Invitrogen; A1049101) supplemented with 10% FBS (Invitrogen; 10270-106), penicillin/streptomycin (Sigma; P4083) at density between 0.5 x 10^6^ cells/ml. PCM6 cells were cultured in McCoy’s 5A modified medium (Gibco;16600082), supplemented with 20% FBS (Invitrogen; 10270-106), penicillin/streptomycin (Sigma; P4083) and 30 U/ml recombinant IL-6 (PeproTech; 200-06). Media were changed every 3 days and replated typically every 5 days the proper dilution was by cell count every time. The lentiviral transfections were performed as described for PE2 cells. For knocking-out the mutant DIS3 allele CRISPR/Cas9 methodology by introducing a truncating eGFP signal as described previoslly^57^. The gRNA sequence (TCGGATGCGTTTATATACGG) was cloned into the pRS1208 – pSpCas9(BB)-2A-Puro (PX459) plasmid (Addgene plasmid #48139). The mCerulean sequence was replaced by *EGPF* using SLIC protocol^56^ in the mCerulean_p2a_neo C-Terminal Fusion (Plasmid 1A)^57^. The homology donor sequence was produced by PCR using primers: GTAATTGTGCTACAAACAGTTCTTCAAGAAGTGAGAAATCGCAGTGCCCCCGTATATggagctggtgcaggcg and TAGTGAAAGTATAGAAATGCTTCTCTTGGTTATTAGTCACATCTCGGATGCGTTTatgggtggaggcggttca. 10^7^ cells in 200 ul of RPMI 1640 without additives were electroporated with a Gene Pulser at a capacity of 960 μF and 180V, in a single exponential decay pulse. The cell suspention was immediately removed from the cuvette and pipetted into another tube containing 500 μl fresh medium without additives. After centrifugation cells were resuspended in 5ml pre-warmed full medium for further 3 day selection in puromycin and subsequent culture at standard conditions. Cells were analyzed on a CytoFLEX LX (Beckman Coulter) flow cytometer. The proper integration of insert was confirmed using PCR amplification with primers spaning over the insertion site (Insertion1_F: tgtgggccaacgtattcctt; Insertion1_R: tgaacttgtggccgtttacg; Insertion2_F: CTTGAGGACCCTGCCATCAG; Insertion2_R: tgaacttgtggccgtttacg) followed by sanger sequencing.

### CellTrace Proliferation Assay

CellTrace™ Yellow Proliferation kit was used according to the manufacture’s instructions. Briefly: Cells were stained by adding 1 μl of CellTrace™ Yellow (Invitrogen, Cat #C34567) stock solution in DMSO per each ml of 10^6^/ml cell suspension in PBS (5 μM working concentration), and subsequently incubated for 40 minutes at room temperature, while being protected from light. Afterward, five times the original staining volume of culture medium, containing FBS, was added to the cells and incubated for 5 minutes. The cells were then pelleted and resuspended in a fresh pre-warmed complete culture medium. At least 4 h incubation period was allowed before analysis to enable the acetate hydrolysis of the CellTrace™ reagent. Cells were pipetted in technical triplicates, 3×10^5^ per well, into round bottom 96 well plate in 200 μl growth medium respective for the cell line tested. Unstained control cells were parallelly cultured in triplicates. Before cytometric analysis, dead cells were stained with LIVE/DEAD fixable dyes (Invitrogen™) according to manufactures instructions and analized on a CytoFLEX Flow Cytometer (Beckman Coulter). Data were analyzed using FlowJo (v10.7.1) software. To calculate the rate of fluorescent signal dilution autofluorescence, defined as the signal from unstained samples, have been subtracted from the fluorescence of all the stained cells and initial (0 h) values have been divided by ones measured at subsequent days of culture assayed.

### Mouse line generation

Dis3 knock-out (KO) mouse lines with a frame-shift insertion in exon 3 (p.K154EfsX24) in C57BL/6N genetic background, were established using the CRISPR/Cas9 method (gRNA: CATCCGAGTCGCAGCGAAG [TGG]). Briefly, chimeric sgRNAs were synthesized by T7 RNA Polymerase in-vitro transcription and purified by PAGE. Cas9 mRNA was in-vitro transcribed with T7 RNA Polymerase and polyadenylated using *E. coli* Poly(A) polymerase (NEB), and m7Gppp5′N cap was added using Vaccinia Capping System (NEB). To induce superovulation, donor mice were injected with 10 IU of PMSG (Pregnant Mare Serum Gonadotropin; Folligon, Intervet, Netherlands) and after ∼50 hours with 10 IU of hCG (Human Chorionic Gonadotropin; Chorulon, Intervet, Netherlands) and mated immediately after hCG injection. Zygotes were collected from mated females 21–22 h post-hCG injection and microinjected into the cytoplasm using Eppendorf 5242 microinjector (Eppendorf-Netheler-Hinz GmbH) and Eppendorf Femtotips II capillaries with the following CRISPR cocktail: Cas9 mRNA (25 ng/μl) and sgRNA (15 ng/μl). After overnight culture, microinjected embryos at the 2-cell stage were transferred into the oviducts of 0,5-day p.c. pseudo-pregnant females. Pups were genotyped at around 4 weeks by PCR using primers: Dis3KO_F: CCTCCTTAAACTCCAGTGGC and Dis3KO_R: CGCAGCCTGATCCTAATGTT, followed by Sanger sequencing of the amplicon.

### Mice breeding conditions

Mice were bred in the animal house of Faculty of Biology, University of Warsaw, and maintained under conventional conditions in a room with controlled temperature (22 ± 2°C) and humidity (55 ± 10%) under a 12 h light/12 h dark cycle in open polypropylene cages filled with wood chip bedding (Rettenmaier). The environment was enriched with nest material and paper tubes. Mice were fed ad libitum with a standard laboratory diet (Labofeed B, Morawski). Animals were closely followed-up by the animal caretakers and researchers, with regular inspection by a veterinarian, according to the standard health and animal welfare procedures of the local animal facility.

### RNA-seq

Total RNA from collected tissues and primary cultures was isolated using a standard TRIzol protocol followed by ribodepletion with a Ribo-Zero Gold rRNA Removal Kit (Human/Mouse/Rat) according to the manufacturer’s instructions (Epicentre; RZG1224). Strand-specific libraries were prepared using a TruSeq RNA Sample Preparation Kit (Illumina) and the dUTP method. Three independent replicate sample sets were prepared for each condition. Libraries were sequenced using Illumina NextSeq platform in a paired-end mode with different average sequencing depth of 19M reads.

### Bioinformatic analysis

MMRF CoMMpass study whole exome DNA sequencing (WES) reads were mapped to the GRCh38 human reference genome using STAR split read aligner^58^, and mutations in *DIS3* were identified, filtered, and quantified using SAMtools and BCFtools pipeline^59^. Healthy tissue control WES data were mapped and variants that were identified in at least one control sample were defined as germline mutations. Whole-genome DNA sequencing reads were mapped to the GRCh38 human reference genome, using STAR. To identify translocations, significant structural variants were called using SVDetect ^60^ with default settings. Significant translocations were filtered for those known to be commonly initiated in the immunoglobulin heavy chain (IGH) locus and genotype. RNA sequencing reads were mapped using STAR. RNA sequencing samples of patients genotyped based on exome sequencing as being a *DIS3* somatic mutation or a recurrent *DIS3* mutation were compared to RNA samples from patients not possessing such a genotype as well as not carrying a germline *DIS3* mutation or a mutant *DIS3* variant as a minor clone. Reads were counted onto a gene annotation (Gencode v34 basic) using FeatureCount (subread package release 2.0.0)^61^. Differential expression RNA-seq analyses were performed as previously using DESeq2 R package^62^.

### Data, code and materials availability

The sequencing data discussed in this publication have been deposited in NCBI’s Gene Expression Omnibus and are accessible through GEO Series accession number: GSE155631.

Any other data, materials, code and additional information required to reanalyze the data reported in this paper is available from the lead contact upon request

### Ethical issues

All procedures were approved by the First Local Ethical Committee in Warsaw affiliated at the University of Warsaw, Faculty of Biology (approval numbers WAW/092/2016, WAW/177/2016, WAW/642/2018). Housing in animal facilities was performed in conformity with local and European Commission regulations under the control of veterinarians and with the assistance of trained technical personnel

## Supporting information

File S1. Contains supplementary information on the mutations in DIS3 detected in CoMMpass study patients

Dataset S1 Exel file. Contains results of differential gene expression analysis from patients with a non-recurrent DIS3 mutation compared to DIS3 WT

Dataset S2 Exel file. Contains results of differential gene expression analysis from patients with a recurrent DIS3 mutation compared to DIS3 WT pati

Dataset S3 Exel file. Contains results of differential gene expression analysis from DIS3 LOH patients upon the loss of mutant DIS3 variant

Dataset EV4 Exel file. Contains results of differential gene expression analysis from PE2 cells upon the reintroduction of WT DIS3

## Acknowledgments

We thank Justyna Chlebowska, Jakub Gruchota, Kamila Kłosowska-Kosicka, Dorota Adamska, Michał Kamiński, Katarzyna Prokop, Patrycja Kędzierska, Weronika Dudzińska and Monika Kusio-Kobiałka for help with selected experiments, Maria Anna Ciemerych-Litwinienko, Dominika Nowis, Jakub Gołąb, Joanna Kufel, Katarzyna Matylla-Kulińska for attentive readings of the manuscript and all Andrzej Dziembowski lab members for fruitful discussions, and Michael Kuehl for sharing the PE2 cell line. Efforts of the Multiple Myeloma Research Foundation (MMRF) and centers that contributed to the CoMMpass study are also acknowledged.

This work was supported by grant funding from the National Science Centre (NCN) to AD (UMO-2013/10/M/NZ4/00299; UMO-2016/22/A/NZ4/00380), TK (UMO-2019/32/C/NZ2/00558) and SM (UMO-2017/27/B/NZ2/01234). This research was also supported by funding from the European Union’s Horizon 2020 research and innovation program (grant agreement no. 810425).

## Author contributions

TK: conceptualization, all bioinformatic analysis, and experimental work; OG, SM, MS, KS, VL: experimental work; AD: conceptualization, supervision, original draft preparation.

## Competing interests

All authors declare that they have no competing interests.

## Notes

### Competing Interest Statement

The authors have declared no competing interest.

### Summary of Updates

We conducted an extra experiment to reproduce the observed counterselection seen during the progression of the disease in humans using a cellular model. This involved employing CRISPR to knock out the D488N mutant variant of DIS3 in a KMS-27 cell line derived from a patient. In Figure 3, we present the results of a prolonged co-culture of KMS-27 MM cells with the DIS3 D488N knockout, illustrating a gradual increase in the contribution of hemizygous wild-type (WT) cells to the total population, as quantified by GFP fluorescence. Additionally, a significant portion of the paper has undergone rewriting for clarity and coherence.

