## SUPPLEMENTAL DATA for "Somatic DIS3 mutations in Multiple Myeloma arise early in clonal evolution, but are later counterselected due to toxicity"

**Kuliński et al**

### SUPPLEMENTAL DATA

- **Figure S1**
- **Figure S2**
- **Figure S3**
- **Figure S4**
- **Figure S5**
- **Description of Additional Information and Supplementary Datasets**

### Supplementary Figures

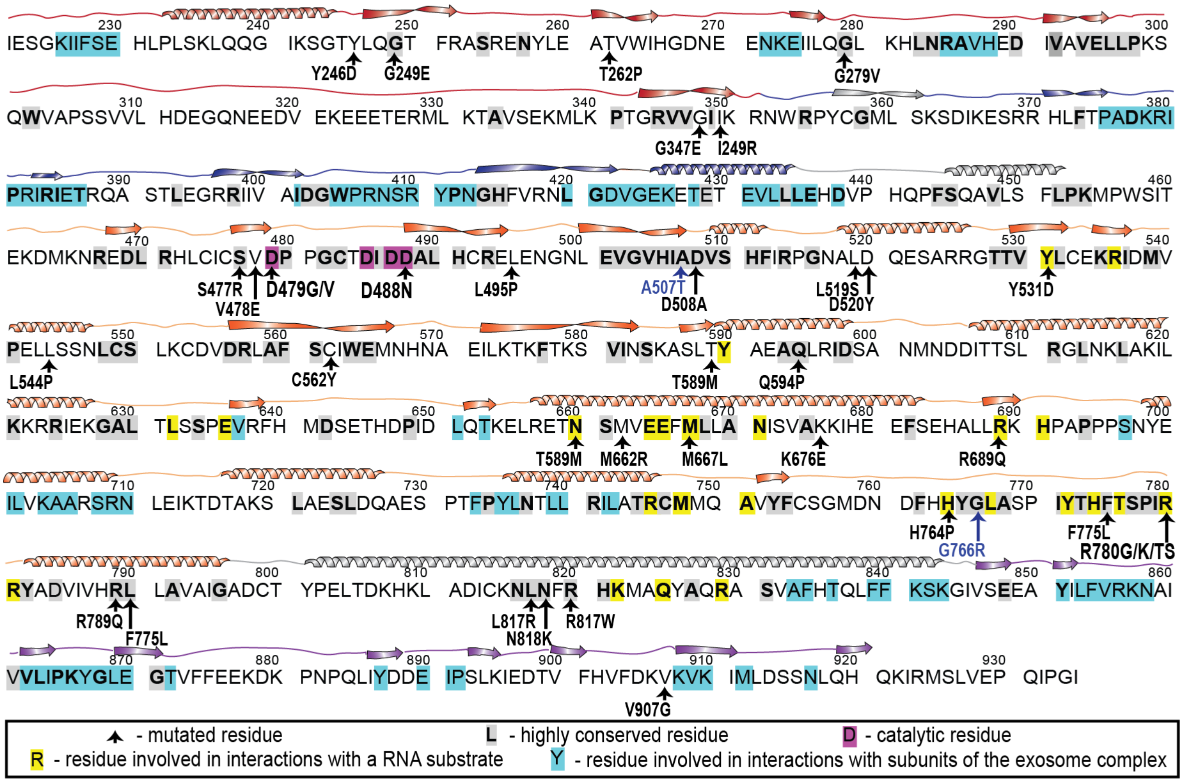


**Figure S1.** Distribution of DIS3 mutations over the amino acid sequence. Highly conserved residues are bolded and highlighted. Catalytic residues, residues that interact with the RNA substrate, and other subunits of the nuclear exosome are in magenta, yellow, and cyan, respectively, according to (Lorentzen *et al*, 2008). A total of 55% of all somatic mutations are in highly conserved residues, and 18% are in residues that interact with RNA substrate. Neither mutations in residues that are responsible for the DIS3 interaction with other components of the exosome complex nor nonsense mutations were identified.

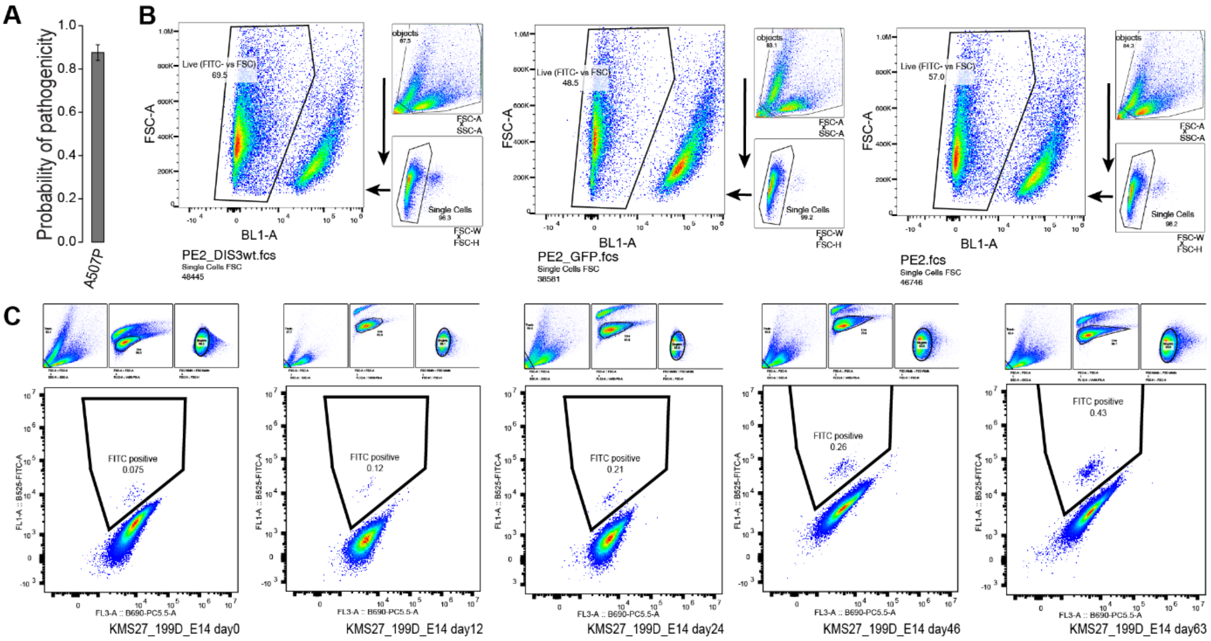


**Figure S2.** Wild type DIS3 reconstitution increases cell viability and proliferation in MM derived cell lines. (A) Prediction of pathogenicity of A507P DIS3 mutation by PON-P2 (B) Gating strategy and representative examples of flow cytometry analyses of PE2 cell viability experiments. (C) Gating strategy and representative examples of flow cytometry analyses of KMS-27 GFP+ cells, with a deleted mutant DIS3 allele outgrowing the population of cells.

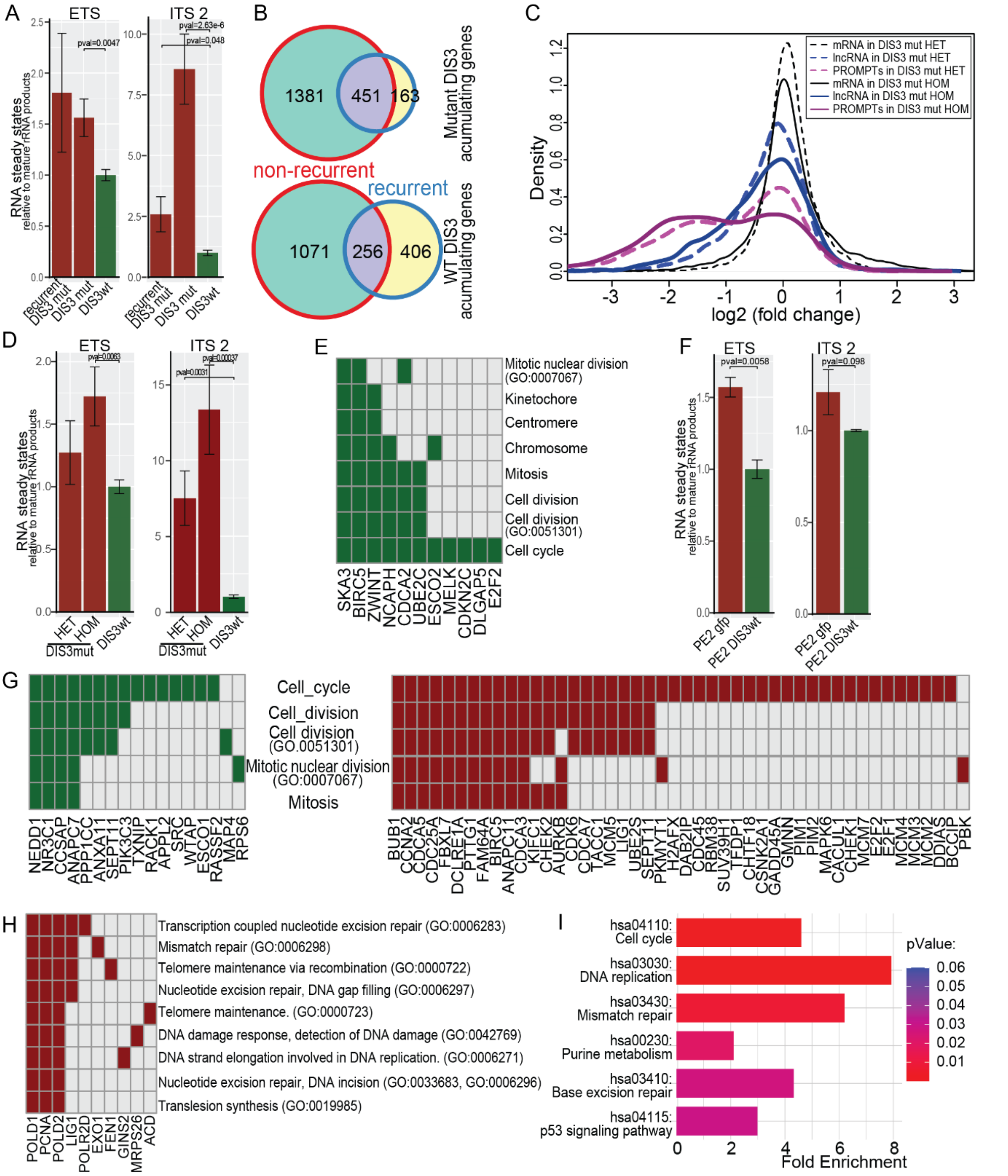


**Figure S3.** Multiple myeloma DIS3 mutations affect its enzymatic activity. (A) Steady-state levels of rRNA processing byproducts 5′ ETS (External Transcribed Sequence) and ITS2 (Internal Transcribed Sequence) normalized to mature rRNA and quantified in RNA-seq data from MM CoMMpass patients, binned according to their DIS3 genotype and shown relative to WT DIS3 patients. Error bars represent the standard error of the mean. (B) The overlap of significantly deregulated genes (Adjusted p<0.01) in somatic non-recurrent and recurrent mutated patients differential expression annalisis. (C) Comparison of the dysregulation of PROMPTs in RNA-seq samples from patients with non-recurrent DIS3 mutations in heterozygous and homozygous states. (D) Comparison of steady-state levels of 5′ ETS and ITS2, normalized to mature rRNA and quantified in RNA-seq data from MM CoMMpass patients with non-recurrent DIS3 mutations, binned according to their zygosity. Error bars represent the standard error of the mean. (E) Cluster of mitosis-related GO terms overrepresented among genes significantly upregulated in samples from DIS3 mutant LOH patients. (F) Levels of rRNA processing byproducts 5' ETS and ITS2 in PE2 cells. Error bars represent the standard deviation. (G) Cluster of mitosis related GO terms overrepresented among genes significantly upregulated in PE2 cells expressing WT (green) or mutant (red) DIS3. (H) Cluster of DNA damage related GO terms overrepresented among genes significantly upregulated in PE2 cells expressing mutant DIS3. (I) Significant enrichment of KEGG pathways among genes significantly upregulated in PE2 cells expressing mutant DIS3.

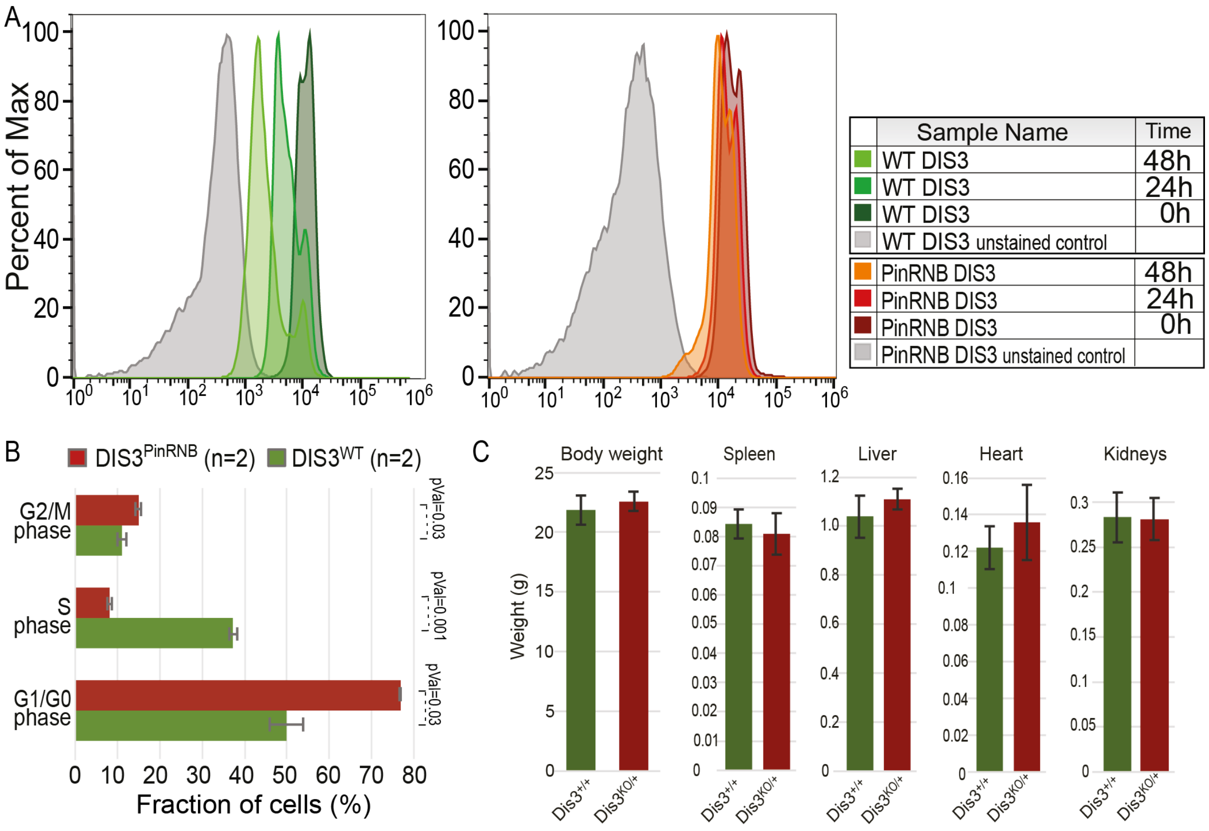


**Figure S4.** DIS3$^{PinRNB}$ allele leads to significant proliferation inhibition. (A) CFSE proliferation assay. (B) Quantification of the faction of cells at different stages of the cell cycle 3 days after induction of DIS3$^{PinRNB}$ expression in 239HEK cells. (C) Weight of 36- to 37-week-old female mice and their major organs (n=6).

**Description of Additional Information and Supplementary Datasets**

**File S1.** Contains supplementary information on the mutations in DIS3 detected in CoMMpass study patients.

1. Table S1 List of DIS3 somatic missense mutations identified in CoMMpass study patients.
2. Table S2 List of DIS3 germline missense mutations identified in CoMMpass study patients.

**Dataset S1** Exel file. Contains results of differential gene expression analysis from patients with a non-recurrent DIS3 mutation compared to DIS3 WT patients

**Dataset S2** Exel file. Contains results of differential gene expression analysis from patients with a recurrent DIS3 mutation compared to DIS3 WT patients

**Dataset S3** Exel file. Contains results of differential gene expression analysis from recurrent DIS3 mutant LOH patients upon the loss of mutant DIS3 variant

**Dataset S4** Exel file. Contains results of differential gene expression analysis from PE2 cells upon the reintroduction of WT DIS3

**Data, code and materials availability**

The sequencing data discussed in this publication have been deposited in NCBI's Gene Expression Omnibus and are accessible through GEO Series accession number: GSE155631.

Any other data, materials and additional information required to reanalyze the data reported in this paper is available from the lead contact upon request.
